# Annotating Gene Ontology terms for protein sequences with the Transformer model

**DOI:** 10.1101/2020.01.31.929604

**Authors:** Dat Duong, Lisa Gai, Ankith Uppunda, Don Le, Eleazar Eskin, Jingyi Jessica Li, Kai-Wei Chang

## Abstract

Predicting functions for novel amino acid sequences is a long-standing research problem. The Uniprot database which contains protein sequences annotated with Gene Ontology (GO) terms, is one commonly used training dataset for this problem. Predicting protein functions can then be viewed as a multi-label classification problem where the input is an amino acid sequence and the output is a set of GO terms. Recently, deep convolutional neural network (CNN) models have been introduced to annotate GO terms for protein sequences. However, the CNN architecture can only model close-range interactions between amino acids in a sequence. In this paper, first, we build a novel GO annotation model based on the Transformer neural network. Unlike the CNN architecture, the Transformer models all pairwise interactions for the amino acids within a sequence, and so can capture more relevant information from the sequences. Indeed, we show that our adaptation of Transformer yields higher classification accuracy when compared to the recent CNN-based method DeepGO. Second, we modify our model to take motifs in the protein sequences found by BLAST as additional input features. Our strategy is different from other ensemble approaches that average the outcomes of BLAST-based and machine learning predictors. Third, we integrate into our Transformer the metadata about the protein sequences such as 3D structure and protein-protein interaction (PPI) data. We show that such information can greatly improve the prediction accuracy, especially for rare GO labels.

## 1 Introduction

Predicting protein functions is an important task in computational biology. With the cost of sequencing continuing to decrease, the gap between the numbers of labeled and unlabeled sequences continues to grow [18]. Protein functions are described by Gene Ontology (GO) terms [16]. Predicting protein functions is a multi-label classification problem where the input is an amino acid sequence and the output is a set of GO terms. GO terms are organized into a hierarchical tree, where generic terms (e.g. cellular anatomical entity) are parents of specific terms (e.g. perforation plate). Due to this tree structure, if a GO term is assigned to a protein, then all its ancestors are also assigned to this same protein. When analyzing only the amino acid sequence data to predict protein functions, there are two major trends. The first trend relies on string-matching models like Basic Local Alignment Search Tool (BLAST) to match the unknown sequence with labeled proteins in the database [11]. Recently, Zhang <u>et al.</u> [18] combined BLAST with Position-Specific Iterative Basic Local Alignment Search Tool (PSI-BLAST) to retrieve even more labeled proteins which are possibly related to the unknown sequence. The key idea behind BLAST methods is to retrieve proteins that resemble the unknown sequence; most likely, these retrieved proteins will contain similar evolutionarily conserved regions and motifs (e.g. kinase domain) that match well with the unknown amino acid sequence. Then, all GO labels assigned to these retrieved proteins are assigned to the unknown sequence [18]. BLAST methods do not explicitly estimate how much a specific motif affects the predicted probability for a GO label.

The second trend transforms the amino acid sequences into features, then applies classification methods to these features. For example, DeepGO converts an amino acid sequence into a string of k-mers, where each k-mer is represented by a vector so that the amino acid sequence is represented as a matrix *m* [8]. Given *m*, the next objective is to find a function *f* (*m, g*) that returns the correct assignment for the label *g*. Neural network models like DeepGO and related work on DNA sequences use the convolutional neural network (CNN) as the key component for this function *f* [8, 19].

In principle, neural network methods are different from BLAST methods; for a specific GO label, a neural network model tries to extract amino acid patterns in the train data that are most correlated with label *g*. When a test sample contains these amino acid patterns, then *f* will return a high predicted probability for *g* given this test sample. To obtain higher accuracy, one may need a very complex neural network model that can extract more information from the training samples. However, due to the complex nature of deep neural networks, it is difficult to say how such patterns, if found, are being analyzed by the models.

In this paper, first, we introduce a novel method GOAT: **GO a**nnotation based on the **T**ransformer framework [17]. In the context of protein sequences, the Transformer architecture is better than a convolutional neural network, because the Transformer models all the pairwise interactions for the amino acids in the same sequence and can possibly capture meaningful long-range relationship among these amino acids.

Second, we reconcile the gap between BLAST and neural network models. As explained, BLAST models retrieve sequences with conserved domains and motifs similar to the unlabeled protein, but do not apply machine learning techniques on these motifs to predict the GO labels. Neural network models want to correlate these motifs and the specific labels via the function *f*, but are not guaranteed to discover such amino acid patterns, especially when the annotated datasets are sparse. For this reason, we identify motifs in the protein sequences via tools like PROSITE [14], and then use this information as extra features for the label classification.

Third, we evaluate how protein metadata such as 3D structures and protein-protein interaction (PPI) network data can affect the prediction. Our work focuses on neural network models, whereas previous work such as Zhang <u>et al.</u> [18] focused on BLAST-based methods.

In the following sections, we describe two baseline methods and the Transformer architecture in our software GOAT. Next, we introduce three types of metadata as extra features for GOAT: Domain (e.g. motifs), 3D-structure, and PPI network information. We find that in the absence of metadata, GOAT is better than DeepGO [8], especially for rare GO terms. Additionally, GOAT with protein metadata obtains higher recall rates than the BLAST-based model in [18]. Our ablation study shows that motif data and 3D folding information are meaningful factors, but PPI network is the most important. We hope that our software GOAT will serve as a meaningful baseline for future research on neural network models for GO annotations. GOAT is available at https://github.com/datduong/GOAnnotationTransformer.

## 2 Method

### 2.1 BLAST and PSI-BLAST

We describe our first baseline which is a very competitive BLAST-based method by Zhang <u>et al.</u> [18]. Zhang <u>et al.</u> [18] consider several best matched proteins from BLAST and additional sequences attained by PSI-BLAST when predicting annotations. The authors [18] refer to their BLAST and PSI-BLAST method as the sequence-based module of their software MetaGO. Here, we refer to it as MetaGO_BLAST_, and briefly describe this method. Let *N*^blast^ and *N* ^psiblast^ be the number of sequences in the train data retrieved by BLAST and PSI-BLAST that best match the unknown protein, and *q* be a GO label. Next define *N*^blast^(*q*) and *N* ^psiblast^(*q*) as the number of sequences having the label *q* in *N*^blast^ and *N* ^psiblast^. Let 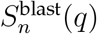 and 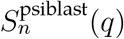 be the sequence-similarity scores of the *n*^*th*^ retrieved sequence in *N*^blast^(*q*) and *N* ^psiblast^(*q*) which contains the GO label *q*. The assigned prediction probability for label *q* with respect to the unknown query sequence is

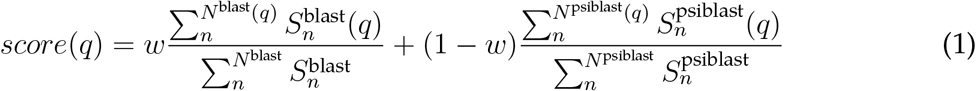

where 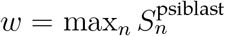 so that BLAST has a stronger weight when very close homologs are found [18]. In other words, the prediction scores for the GO labels are scaled by how similar the query protein is to the known sequences.

### 2.2 Convolutional neural network

We describe our second baseline DeepGO Kulmanov <u>et al.</u> [8] which is built from the convolution neural network (CNN) architecture. We consider the DeepGO version which does not correct for consistent predicted probabilities. Consistency is defined as the fact that if a GO label is assigned to a protein then all its ancestors must also be assigned to the same protein. This consistency is corrected for each GO label, where the final prediction is the maximum of its own predicted probability and the probabilities of its descendants. In other words, in DeepGO when a descendant of a specific label has high predicted probability then this label will also have high predicted probability. There are other types of correction [10] which were not compared in DeepGO. We believe this research direction requires its own analysis. Moreover, consistency correction relies on the per-term prediction accuracy. For this reason, we focus comparing GOAT to only the basic DeepGO.

DeepGO converts an amino acid sequence, for example *p* = MARS · · ·, into a list of overlapping 3-mers, e.g. MAR ARS · · ·. Each 3-mer is represented by a vector of in ℝ^128^, so that *p* is represented by a matrix *E*_*p*_ ∊ ℝ^128*×*(*L−*2)^ where *L* is the sequence length. A 1D-convolution layer, 1D-maxpooling and Flatten are then applied to *E*_*p*_, so that we have *v*_*p*_ = flatten(maxpool (conv1d(*E*_*p*_)) as the vector representing this k-mer sequence. To predict a GO label *i*, DeepGO fits a logistic regression layer sigmoid 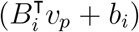 with the binary cross entropy as the objective loss function.

To integrate metadata about the protein (e.g. PPI network), DeepGO concatenates this information into *v*_*p*_ before sending it through the logistic layer. For example, in the original paper, Kulmanov <u>et al.</u> [8] convert proteins from a PPI network into vectors. Let *c*_*p*_ ∊ ℝ^*d*^ be the vector representing protein *p* in this PPI network; then the classification layer becomes sigmoid 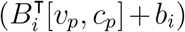 where [*v*_*p*_, *c*_*p*_] is the concatenation of the two vectors. We emphasize that the vector *c*_*p*_ does not have to be from the PPI network; in the results, we evaluate the DeepGO model with vectors representing 3D structures of proteins (the outcome of this experiment is not shown in table but explained in result section).

### 2.3 GO annotation method with Transformer

We introduce our novel **GO a**nnotation method with **T**ransformer (GOAT), available at https://github.com/datduong/GOAnnotationTransformer. GOAT takes as an input a sequence of amino acids and GO labels. For illustration purposes, consider the protein MAP Kinase-activated Protein Kinase 5 (UniProtKB O54992), which we will refer to by its id O54992 for brevity. Let *L* denote the sequence length, and let *g*_1_ · · · *g*_*G*_ denote the names of the labels to be predicted, where *G* is the number of labels to be predicted. Our input will be the string *MSEDS · · · LPHEPQ g*_1_ · · · *g*_*G*_ of total length *L* + *G*. Next, define *E* as the embeddings for the amino acids, for example *E*_*M*_ and *E*_*S*_ are the vectors representing the amino acids *M* and *S* respectively.

Let *E*^*G*^ be the embeddings for the GO labels, so that 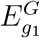 and 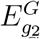 are the vectors representing the first a nd s econd GO label *g*_1_ and *g*_2_ respectively. *E*^*G*^ is a nalogous to Word2vec embedding; except in this case, instead of having a vector for each word in a corpus, we will have a vector for each GO label in the train and test datasets. In this paper, we set the vectors represent the amino acids and GO labels to be in the same dimension; that is, *E*_*M*_ and 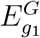 are vectors of the same size. To reduce the number of parameters, for the GO labels, we fix *E*^*G*^ as the definition-based GO embeddings from [6] instead of setting it as a trainable parameter.

We add a position vector *P*_*j*_ to the *j*^*th*^ amino acid in the sequence; for example, the first and second amino acid *M* and *S* will have the following two vectors, *E*_*M*_ + *P*_1_ and *E*_*S*_ + *P*_2_. We observe that the position embedding makes sense for amino acids so that the same amino acid appearing at different locations will be treated differently. However, position embedding does not apply to GO labels; that is, the ordering of the labels should not affect the prediction outcome. For this reason, we do not add position embedding to the GO labels.

Next, we introduce the region-type embeddings *R* to highlight the fact that some amino acids belong to known motifs. Again consider the protein O54992, which is 473 residues long and contains two key regions: a kinase motif at position 22-304 and a coiled coil domain at position 409-440. In this case the 25^*th*^ amino acid is *T* and is inside the kinase motif; for this reason, it will have the vector *E*_*T*_ + *P*_25_ + *R*_*kinase*_. Likewise, the 410^*th*^ amino acid is *N* and will have the vector *E*_*N*_ + *P*_410_ + *R*_coiled coil_. Amino acids outside any key regions will not have region embedding added to them. Motifs can be found by using PROSITE; fortunately, many labeled sequences in Uniprot already have this information [5, 14].

We now describe how the Transformer architecture in GOAT analyzes the input sequence. The original Transformer has 12 layers of encoders and each layer has 12 independent identical units (referred to as heads in the original paper) [17]. To keep our software manageable for all users, in this paper, we use only one head and so we will exclude description of head. We will use 12 layers. The first layer takes as arguments the vectors representing the input string. Here we simplify the notation, let *w*_*j*_ be the vector representing the *j*^*th*^ element in the input string. For protein O54992 of length *L* = 473 and *G* number of GO labels, from the input string *MSEDS · · · LPHEPQ g*_1_ · · · *g*_*G*_, we will have for example *w*_25_ = *E*_*T*_ + *P*_25_ + *R*_kinase_, and *w*_474_ = *E*_*g1*_. At the first layer, we have

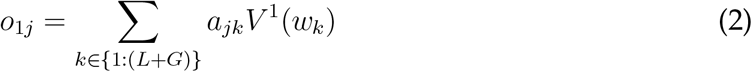

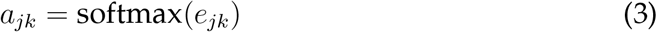

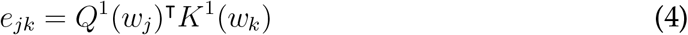

*V* ^1^, *Q*^1^, *K*^1^ are transformation functions for layer 1. *o*_1*j*_ is a weighted sum of the transformed vectors representing itself and the other entities in the input string. *o*_1*j*_ is then transformed as 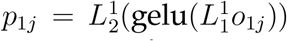 where 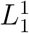 and 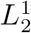 are two linear transformations with the gelu activation function in between. The final output of Layer 1 is *h*_1*j*_ = LayerNorm(*p*_1*j*_ + *o*_1*j*_). Loosely speaking, the first layer computes all pairwise interactions of *w*_*j*_ and *w*_*k*_ for all *k*, where the attention *a*_*jk*_ in Eq. 3 indicates how much *w*_*k*_ contributes toward *w*_*j*_.

The second layer takes the output of the first layer as its input, so that we have for any layer *i*

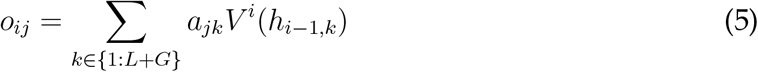

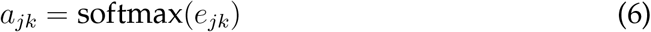

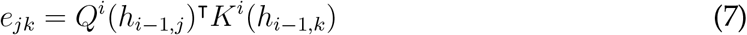

where *V*^*i*^, *Q*^*i*^, *K*^*i*^ are transformation functions for layer *i*. This layer *i* will have its own linear transformations 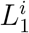 and 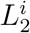 to transform *o*_*ij*_. Again, loosely speaking, layer *i* computes all pairwise interaction for the output from the previous layer *i* − 1.

At layer 12, we focus only on the output *h*_12,*k*_ corresponding to the GO labels. Let us denote 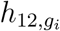 as the final output for the term *g*_*i*_. We use a single linear classifier softmax 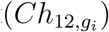 to return the presence and absence probability of *g*_*i*_ for the input protein. The same transformation *C* is applied to all labels, so that a set *S* of GO terms having similar 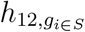 will have similar predictions.

At each label *g*_*i*_, the output 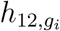 encapsulates all the information from the amino acids and from all the other labels, so that values which affect 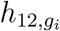 will also affect the output 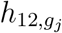 at another label *g*_*j*_. Intuitively, with this fact and the fact that all labels share the same classifier *C*, the prediction at *g*_*i*_ and *g*_*j*_ are to some degree correlated. We suspect that Transformer can model co-occurrences of labels. To validate this, from the T-SNE plot of the vectors 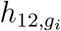 in the result section, we observe that a label and its ancestors will be nearby, even when their initial definition embeddings 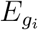 in [6] are dissimilar, as is the case when a term and its distant ancestors can have dissimilar definitions.

When there are metadata for the proteins, such as their embedding *c*_*p*_ from a PPI network, we can concatenate such embeddings into 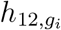 as in DeepGO. Next, we use a two-layer fully connected classifier, softmax 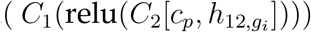 to return the presence and absence probability of *g*_*i*_, where *C*_1_ and *C*_2_ are shared for all the labels.

## 3 Results

We present three main results in the following sections. First, when using amino acid sequence data alone as input, we show that our adaptation of Transformer in GOAT is better than the CNN-based DeepGO at predicting protein functions. We will use the name GOAT_BASE_ and DeepGO_BASE_ to indicate the base implementation in both models, and ignore the subscripts when the context is clear. Second, we show that GOAT obtains higher classification accuracy when it takes as extra inputs the motif information from the protein sequences. We use the name GOAT_MOTIF_ for this version of GOAT. We also observe how the Transformer architecture in GOAT analyzes the motif information when it predicts GO labels for an amino acid sequence. Third, because motif information is a key input of GOAT, we integrate 3D-structure and PPI network metadata about the proteins on top of our GOAT_MOTIF_ to obtain even better prediction outcome. We use the name GOAT_MOTIF,3D_, GOAT_MOTIF,PPI_, GOAT_MOTIF,3D,PPI_ to indicate joint inputs in GOAT.

We use the datasets from the original DeepGO paper, which contain one dataset each for the BP, MF, and CC ontologies. We remove proteins without any GO annotations. The BP, MF and CC datasets have 27279, 18894, and 26660 train proteins, and 9096, 6305, and 8886 test proteins, respectively. In the DeepGO datasets, BP, MF and CC labels annotating fewer than 250, 50, and 50 proteins are removed. The ancestors of all the terms annotating one protein are also added into the ground truth label set. In total, the numbers of BP, MF and CC terms in the label sets are 932, 589, and 439 respectively.

We measure the per-label accuracy using Macro and Micro AUC which are the un-weighted and weighted averages of the AUC at each label, respectively. We are more interested in the accuracy for rare labels because these labels are closer to the true functions of the proteins. Rare labels affect Macro AUC more than Micro AUC; however a high Macro AUC does not always guarantee that rare labels are correctly classified. For example, the BP term *protein glycosylation* is more precise to a protein function than its ancestors *metabolic process* and *cellular process*. However, by design, DeepGO datasets include all the ancestors for a label. The label *protein glycosylation*, *metabolic process* and now occur 254, 12455 and 19232 times in the data, respectively. If *metabolic process* and *cellular process* are correctly predicted, then Macro AUC is high but does not imply that we can properly predict *protein glycosylation*.

For this reason, to evaluate the models, we also compute recall-at-k (R@k) which measures the per-protein accuracy. In practice, a classifier would return *k* of most probable labels for an unknown sequence, which a curator can then review. These *k* labels are referred to as *top-k labels* because they are the *k* labels having the highest predicted probabilities for an input sequence. For one protein, R@k measures the fraction of correct labels retrieved among the top-k labels. We report the final R@k which is the average R@k over all test samples evaluated with respect to the label set of interest. We evaluate R@k on the entire label set and also on the sets of rare labels.

### 3.1 Base GOAT

To compare the base architectures in GOAT and DeepGO, we fit these two models on only amino acid sequences without any extra information such as motifs or PPI network data. For DeepGO, we use the same hyperparameters as the original paper [8]. For the Transformer parameters in GOAT, we set input embedding at size 256 (that is *E*^*G*^ ∊ ℝ^932*×*256^ for MF), and intermediate vectors at size 512. To keep our software GOAT manageable for all users, we implement Transformer with only one head and 12 layers. We initialize the GO embedding *E*^*G*^ as the pre-trained embedding BERT_LAYER12_ in [6] which transforms the GO definitions into vectors, where GOs having related definitions will have comparable vectors. Using pre-trained GO embeddings reduces the number of parameters in the Transformer, which can also reduce overfitting, use less GPU memory, and decrease run time. We train Transformer on a single GTX 1080 Ti with 11GB memory for all three datasets.

In BP, MF and CC data, GOAT_BASE_ exceeds DeepGO_BASE_ in Macro and Micro AUC, indicating that the base implementation of GOAT attains better per-label accuracy (Table 1 row 3 and 5). To estimate per-protein accuracy, we select R@50, R@30, and R@30 for the BP, MF and CC data, respectively (approximately 5% of total label size). On the whole data, GOAT increases recall by a small amount. Recall for the entire label set can be affected by common GO terms which are often easier to classify compared to rare labels. As discussed in the previous section, accurately predicting rare terms is more important for the proteins, because rarer labels describe more detailed biological events which reflect the true properties of the unknown proteins.

**Table 1:** Macro AUC, Micro AUC and Recall-at-k (R@k) are evaluated on the entire 932 BP, 589 MF, and 439 CC labels in the original DeepGO data. *k* value in R@k corresponds to about 5% of total label size in each dataset.

|  |  | BP |  |  | MF |  |  | CC |  |  |
| --- | --- | --- | --- | --- | --- | --- | --- | --- | --- | --- |
|  |  | Macro | Micro | R@50 | Macro | Micro | R@30 | Macro | Micro | R@30 |
| **BLAST Psi-BLAST** |  |  |  |  |  |  |  |  |  |  |
| 1 | Evalue10 | 65.99 | 81.34 | 48.02 | 76.51 | 87.12 | 69.75 | 64.17 | 89.36 | 77.24 |
| 2 | Evalue100 | 66.61 | 84.41 | 48.84 | 78.67 | 90.34 | 68.06 | 65.62 | 92.54 | 80.23 |
| **DeepGO** |  |  |  |  |  |  |  |  |  |  |
| 3 | BASE | 62.60 | 82.17 | 46.13 | 73.90 | 87.49 | 59.63 | 67.12 | 93.07 | 81.47 |
| 4 | +PPI | 82.16 | 90.31 | 56.97 | 84.97 | 92.40 | 70.19 | 87.51 | 96.76 | **87.98** |
| **GOAT** |  |  |  |  |  |  |  |  |  |  |
| 5 | BASE | 67.69 | 84.46 | 47.40 | 78.67 | 89.43 | 60.46 | 74.94 | 93.95 | 82.98 |
| 6 | +MOTIF | 71.04 | 85.64 | 48.65 | 82.53 | 91.12 | 66.19 | 77.62 | 94.51 | 83.01 |
| 7 | +MOTIF+3D | 72.62 | 86.09 | 50.76 | 85.38 | 92.72 | 69.23 | 79.57 | 94.95 | 84.22 |
| 8 | +MOTIF+PPI | **83.68** | **90.49** | **57.81** | **88.92** | **93.98** | **72.16** | **90.76** | **97.23** | 87.95 |

Table 2 shows the R@k for 232 BP, 143 MF, and 110 CC rare labels which appear below the 25^*th*^ quantile occurrence frequency in the label sets. For recall rates to make sense in this case, we compute recall rates for the proteins annotated by at least one of these rare labels. In the test data, GOAT now has a noticeable improvement over DeepGO, especially for larger sets of top-k labels (Table 2 row 3 and 5). This evaluation indicates that our adaptation of the Transformer framework in GOAT can extract more useful information from the amino acid sequences, and thus obtain higher prediction accuracy.

**Table 2:** Recall-at-k (R@k) are evaluated on the most uncommon 232 BP, 143 MF, and 110 CC labels from the entire set of size 932 BP, 589 MF, and 439 CC labels in the original DeepGO data.

|  |  | BP |  |  | MF |  |  | CC |  |  |
| --- | --- | --- | --- | --- | --- | --- | --- | --- | --- | --- |
|  |  | R@10 | R@20 | R@50 | R@5 | R@10 | R@30 | R@5 | R@10 | R@30 |
| **BLAST Psi-BLAST** |  |  |  |  |  |  |  |  |  |  |
| 1 | Evalue10 | 25.13 | 32.59 | 47.33 | 44.87 | 51.11 | 57.51 | 25.09 | 30.91 | 49.27 |
| 2 | Evalue100 | 21.79 | 29.88 | 46.29 | 42.31 | 48.37 | 60.98 | 21.91 | 29.72 | 50.47 |
| **DeepGO** |  |  |  |  |  |  |  |  |  |  |
| 3 | BASE | 11.88 | 20.21 | 39.75 | 17.42 | 27.07 | 50.47 | 13.28 | 24.57 | 48.56 |
| 4 | +PPI | **45.62** | **57.78** | 74.55 | 41.96 | 52.85 | 71.98 | **53.14** | **64.85** | 82.58 |
| **GOAT** |  |  |  |  |  |  |  |  |  |  |
| 5 | BASE | 13.49 | 23.32 | 44.54 | 18.80 | 30.95 | 58.87 | 17.89 | 26.47 | 51.21 |
| 6 | +MOTIF | 17.27 | 27.75 | 48.11 | 30.45 | 41.72 | 64.76 | 19.06 | 31.03 | 58.94 |
| 7 | +MOTIF+3D | 23.66 | 34.27 | 55.06 | 36.19 | 47.90 | 72.23 | 27.36 | 39.72 | 67.69 |
| 8 | +MOTIF+PPI | 42.84 | 56.01 | **74.78** | **45.35** | **59.49** | **82.67** | 48.04 | 62.14 | **84.42** |

We next evaluate whether our adaptation of Transformer can learn the co-occurrences of labels. Duong <u>et al.</u> [6] noticed in their GO embedding that when one of the child-parent GO labels describes very broad biological events (e.g. low IC), then their vector representations may be far apart. This fact implies that for Transformer to work well, to some degree it must learn the co-occurrences of labels and adjust *E*^*G*^ so that any two related GO labels (regardless of their frequencies in the train data, IC values and distance to roots) will have comparable vectors. To observe that Transformer can implicitly learn label co-occurrences, we compare the T-SNE plots of the input GO embedding *E*^*G*^ and its output 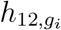 from the Transformer layer 12 which is directly passed into the classification layer.

For every input protein, we have a different value of 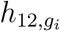 for the same label *g*_*i*_ because 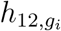 is function of the vector representing the amino acids. We apply our trained Transformer on the test set, and take the average 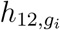 over each input proteins in test data (denoted as 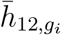). We compute 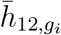 from the test data because these proteins are not seen in training and provide a more realistic evidence. We use 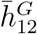 to denote the set of 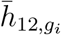 for all *g*_*i*_.

Figure 1 shows the T-SNE plot of *E*^*G*^ for the MF labels in DeepGO dataset; we highlight two terms GO:0008376 (red) and GO:0030291 (blue) and their ancestors. The dot size is scaled by IC values, where smaller size implies lower IC (so the label is more common). Smaller dots tend to cluster well together but large dots do not. For example, consider the term GO:0016740 and its parent GO:0003824 (top right and far left red nodes) which should often co-occur because in our dataset ancestors of an assigned GO label are included as the ground truth labels. For Transformer to work well, it should reposition the 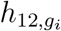 vectors representing GO:0016740 and GO:0003824 closer together.

**Figure 1:**
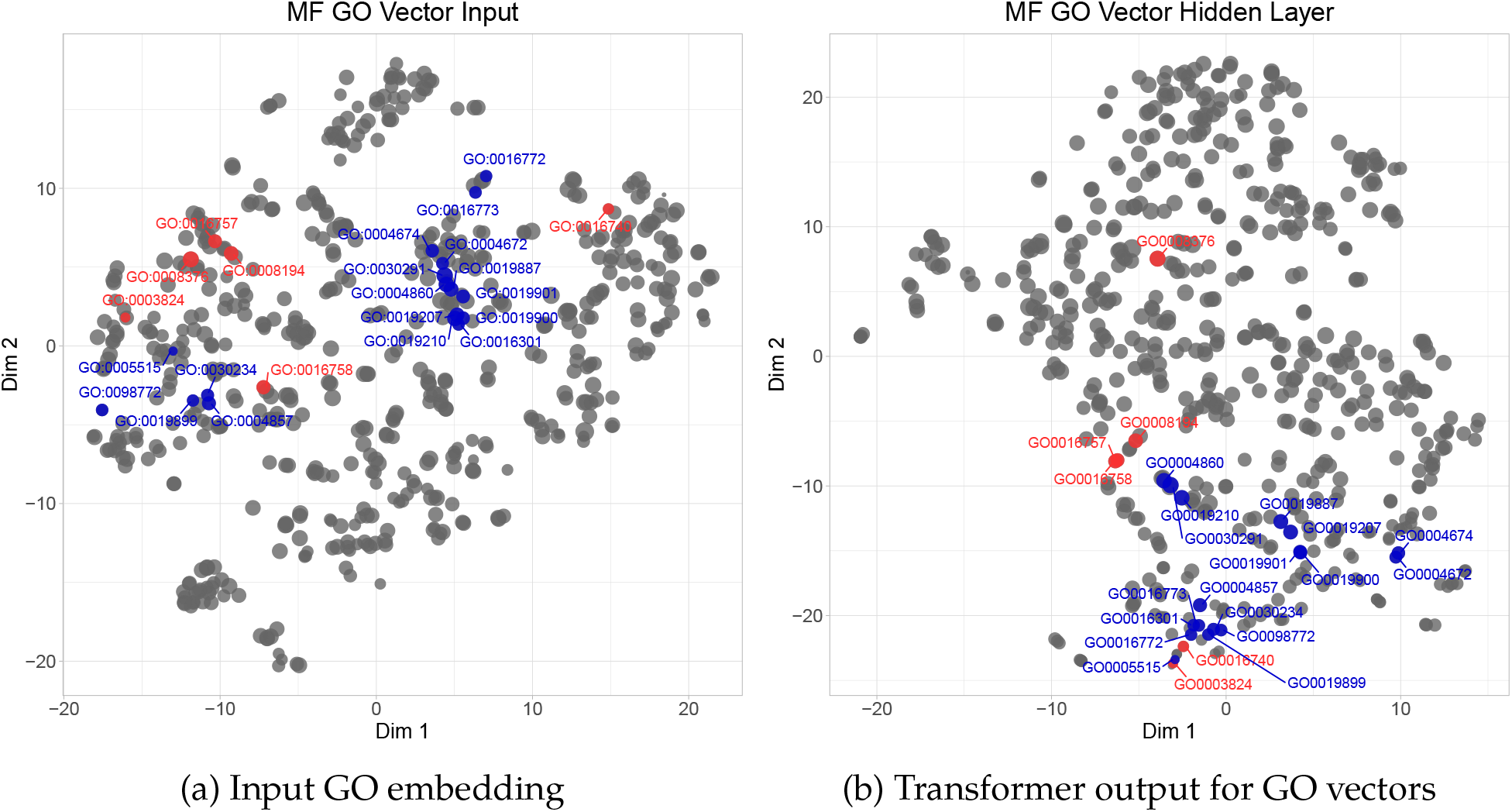
T-SNE of input GO embeddings and their transformed values created by Transformer layer 12.

The T-SNE plot of 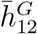 from Transformer shows that the red and blue nodes have clustered more closely compared to *E*^*G*^. The blue dots now gather at the bottom of Fig, as compared to being two separated groups in Fig. The two example cases GO:0016740 and GO:0003824 are now near one another, bottom of Fig. However, the lowest level red node GO:0008376 remains far from its ancestors. The red and blue dots are not yet tightly compacted into two dense clusters, and so there is room for further development. In conclusion, Transformer weakly models the co-occurrences of labels even when such a constraint is not explicitly enforced.

### 3.2 Domains and motifs in amino acid sequences as features

Before the introduction of neural network models, earlier methods often integrated BLAST as a key component. BLAST retrieves annotated proteins in the database that share the same conserved domains or motifs with the unknown sequence, and then assigns the GO labels of these retrieved proteins to the unknown query. Loosely speaking, key domains or motifs shared by the training sequences can then be considered as the key factors in BLAST-based methods. In this section, we evaluate whether neural network models can automatically learn these key patterns from the training sequences, and explain how to introduce these patterns as input features to GOAT.

For fair comparison, we select a very strong BLAST baseline MetaGO_BLAST_ [18] and apply it to the same DeepGO datasets. We emphasize that MetaGO_BLAST_ in [18], which matches multiple related sequences to the query, has much stronger performance than the BLAST baseline used in DeepGO where the authors select only a single best matching sequence [8]. We build the BLAST database from the DeepGO train data. For BLAST and PSI-BLAST, we perform the experiments using both e-value at 10 and 100. Lower e-values leave too many unmatched testing sequences; for example, in the CC dataset at e-value 1, only 7236 out of 8886 test samples match to some sequences in the train data. Moreover, in the context of finding possible protein functions, a higher e-value can allow for higher recall rates.

In BP and MF data, MetaGO_BLAST_ is better than the base DeepGO and GOAT; for example, MetaGO_BLAST_ yields better recall rates on rare labels (Table 1 and 2 row 1–3 and 5). We emphasize that MetaGO_BLAST_ and GOAT_BASE_ have comparable recall on rare MF labels for larger top-k label sets. Arguably, MetaGO_BLAST_ and GOAT retrieve the same number of correct labels, but GOAT makes more spurious predictions. DeepGO however does not come close to MetaGO_BLAST_ on recall rates. The convolutional network in DeepGO is likely not complex enough to learn motifs shared among the sequences with related functions, which BLAST or PSI-BLAST can detect.

When evaluated on the entire CC data, the base DeepGO and Transformer outperform MetaGO_BLAST_. However, for rare CC labels, when compared to MetaGO_BLAST_, Transformer has higher R@30 but lower R@5 and R@10. To some degree, our current GOAT is not yet better than MetaGO_BLAST_.

We had hoped that the complex Transformer architecture in GOAT would have learned key motifs missed by the simpler convolutional neural network in DeepGO, but this is not the case. There are information about the amino acid sequence that we must explicitly specify for GOAT to function better. For this reason, we use the motifs extracted by string-matching methods as inputs to our method GOAT. For example, we can apply PROSITE to scan the protein database for known motifs in a given input sequence [14]. Loosely speaking, we are combining motif-based methods and the Transformer architecture into one pipeline by taking the output of motif-based models and passing them as inputs to our Transformer. Our strategy is different from an ensemble approach that would average the prediction of independent predictors.

In the manually reviewed Uniprot database, each protein already has a Family & Domains section describing its key regions [16]. We then add these region types of the amino acids as input features into our method GOAT. We emphasize that not all domains are meaningful at predicting labels in a certain ontology. For example, Serine/threonine-protein kinase TBK1 (UniProtKB number Q9UHD2) has a kinase domain at position 9-310 (PROSITE annotation rule PRU00159). However, kinase domain involves in a wide range of biological processing at various locations in the cell like metabolism, transcription, cytoskeletal rearrangement and movement, and cell apoptosis and differentiation [7, 13]. Thus, this kinase domain in Q9UHD2 tells us which Molecular Functions are more likely to be assigned, but this domain cannot tell which Biological Processes or Cellular Components Q9UHD2 will have.

For each protein in the original DeepGO datasets, we download its sequence annotation rule (e.g. PROSITE rule) from the 2019 data at https://www.uniprot.org/. We do not consider region types for sequences that have changed in length; otherwise, we consider region types only for portions that have not changed in amino acids composition. Uniprot data divides region types into subgroups. Some subgroups require curated comments and are not truly applicable for analyzing new proteins; for example, the subgroup Domain Non-positional Annotation is not determined by sequence analysis.

We use the following six subgroups which can be found by sequence analysis: Zinc finger (e.g. C2H2-type), Repeat (e.g. AA tandem repeat), Motif (e.g. LXXLL motifs), Compositional bias (e.g. Asp/Glu-rich), Coiled coil (e.g. Leucine-zippers), Domain (e.g. Ser/Thr kinase domain). In the original DeepGO datasets, we found 1629, 1450 and 1655 amino acid region types for the BP, MF and CC train data, respectively. Region types found in test sequences but not seen in the train data are set as zero; effectively we treat these cases as if the region types do not exist. To model the region types in the amino acid sequence, we apply the region-type embedding *R* explained in section; for example, in BP ontology we will have a embedding *R* ∊ ℝ^1629*×*256^. We will use the name MOTIF to denote all types of Domain information in these six subgroups reported by The UniProt Consortium [16].

When evaluated on the entire data, GOAT_MOTIF_ obtains better AUCs and comparable recall rates to MetaGO_BLAST_ (Table 1 row 6). When evaluated on rare labels, our GOAT has subpar recalls when the top-k label set is small; however, for larger top-k label set, especially in MF and CC data, GOAT retrieves more correct labels (Table 2 row 6). This result implies that GOAT_MOTIF_ is a better classifier when the top-k label set is larger. Our result also highlights an important point. Ideally, a neural network should teach itself the patterns associated with certain key protein functions. However, a neural network may fail to learn such key information, and needs these inputs to be explicitly provided. These key domains within a sequence are often easily obtained, and in fact already available in Uniprot.

To observe how the MOTIFS are analyzed by the Transformer architecture in GOAT, we select the human protein Serine/threonine-protein kinase TBK1 (UniProtKB Q9UHD2) in DeepGO test data. As described in section 3.2, we concatenate the amino acid sequence and GO labels into one single input into GOAT. Q9UHD2 is 729 amino acids long, and when predicting MF labels, the input into GOAT is then a string of amino acids and GO labels of length 1318 (from 729 + 589 MF labels). We […something about region-type emb] integrate into the Transformer framework the three key domains in Q9UHD2 derived from sequence analysis methods; these are Protein Kinase at 9-310, Ubiquitin-like at 309-385, and Coiled coil at 407-713. The vertical and horizontal red lines indicate these regions in the attention heatmap (Figure 2).

**Figure 2:**
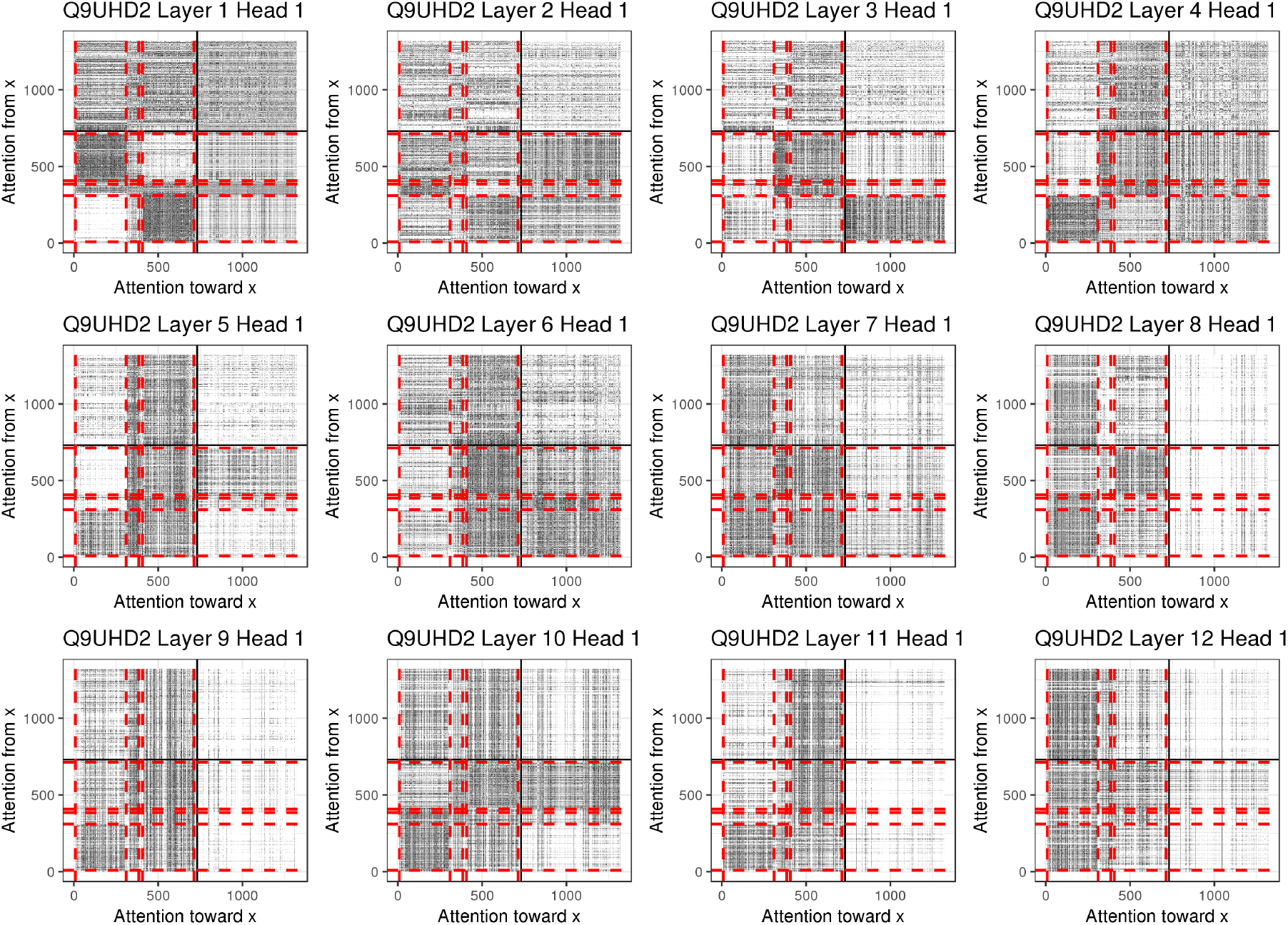
Heatmap of the attention values *α*_*jk*_ in each layer when analyzing the protein kinase TBK1 (UniProtKB Q9UHD2). The three key regions of this sequence (separated by red lines) are explicitly given as inputs to the Transformer model. The first quadrant shows the interactions among the GO labels, the second shows contribution of amino acids toward the GO labels, the third shows interactions of amino acids among themselves, and the fourth shows contribution of GO labels toward the amino acids.

We plot the attention heatmap of *α*_*jk*_ in each layer (Figure 2). *α*_*jk*_ measures how much position *k* contributes toward position *j* (Eq. 3). Each row in the heatmap adds to 1. The heatmap is divided into four quadrants. The first quadrant shows the interactions of GO labels among themselves (e.g. *α*_*jk*_ for *j, k* ∊ [730, 1318]), the second shows contribution of amino acids toward the GO labels, the third shows interactions of amino acids among themselves (e.g. *α*_*jk*_ for *j, k* ∊ [1, 729]), and the fourth shows contribution of GO labels toward the amino acids.

Transformer without region-type embedding has a noisy attention heatmap (Appendix Fig. 3). Transformer with region-type embedding displays meaningful patterns (Fig. 2). For example, layer 1 illustrates the cross interactions between Protein Kinase and Coiled coil domain (black boxes quadrant 3); whereas layer 4 and 8 show interactions within the Protein Kinase and Coiled coil themselves. In layer 12, the final vectors representing GO labels receive more attention from the Protein Kinase than the other regions (top right box quadrant 2). Because these final output vectors are sent to the classification layer, we can assume that the Protein Kinase region contributes more to the label annotation compared the other domains. This observation is consistent with the true molecular functions of Q9UHD2 which are: phosphoprotein binding, protein kinase activity, protein phosphatase binding, and protein serine/threonine kinase activity.

### 3.3 Protein vectors as features

Zhang <u>et al.</u> [18] evaluates how PPI network and 3D structure data affect the annotation accuracy for methods built from BLAST. In this paper, we assess the contributions of these components in the context of a neural network classifier. The amino acid sequences have already been used in neural network models, as described in DeepGO and our GOAT. Any other information about protein must come from some external resources such as the STRING database [15]. For example, one can train a neural network on the STRING database to transform interacting proteins into similar vectors [1, 4]. Integrating these protein vector representations into annotation methods is motivated by the fact that interacting proteins (ideally encoded into similar vectors) should have closely related functions (e.g. found in the same biological processes and cellular locations).

It is only recently that proteins in knowledge graph have been transformed into vectors via neural network models. In this paper, we will not build new models to encode proteins from knowledge graph, and reserve this topic for future work. We will focus on evaluating how much can protein vectors built from external data sources increase the prediction accuracy.

#### 3.3.1 Vectors representing protein 3D structure

We evaluate protein vectors that capture high-level 3D shapes of the proteins (e.g. *α*-helices and *β*-sheets) [3]. [3] applied a 3-layer Bidirectional Long-Short Term Memory (BiLSTM) to encode an amino acid sequence into a matrix. [3] trained their model on the SCOP database and residue-residue contact prediction. The SCOP database describes the major classes of 3D-structures often seen in proteins. Their model was trained on SCOPe ASTRAL 2.06 dataset with 22,408 amino acid sequences, and each training epoch has 100,000 pairs sampled from these 22,408 sequences [3]. From SCOP, Bepler and Berger [3] predict whether two protein sequences have no relationship, class-level relationship, fold-level relationship, superfamily-levelrelationship, or family-level relationship (e.g. label *y* = 0, 1, 2, 3, 4) [3]. For example, two proteins with the same Rossmann-fold structural motif have a class-level relationship (*y* = 1 in this case). Residue-residue contact prediction is applied within the same protein sequence; the objective is to predict whether each pair of positions *i, j* within the same protein are close by (distance less than 8 angstroms) in the 3D structure. Bepler and Berger [3] provide the pre-trained encoder which can return a matrix for any amino acid sequence, even those not used in training. In this paper, we take the mean-pool of this matrix to represent the entire sequence.

Because motif information is a key input of GOAT, we integrate 3D-structure data on top of our GOAT_MOTIF_. We emphasize that 3D-structures and motifs are targeting two different kinds of information; for example, a kinase domain does not strictly entail a specific folding pattern. For this reason, GOAT_MOTIF,3D_ improves upon GOAT_MOTIF_ (Table 1 row 7). More importantly, for a larger top-k label set, GOAT is much better than MetaGO_BLAST_, where our recall rates increase by about 9%, 12%, and 17% in BP, MF and CC data respectively (Table 2 row 7). Our results indicate that GOAT find fewer correct labels when the set of top-k labels is small; however, GOAT will retrieve more correct labels than MetaGO_BLAST_ for larger set of top-k labels.

When we replace the PPI network in DeepGO with the vectors from SCOP in [3], the performance significantly decreases. Macro AUC in DeepGO drops from 82.16 to 66.54 in BP, from 84.97 to 77.78 in MF, and from 87.51 to 68.93 in CC data. This decrement is anticipated as we will argue in the next section, that protein interaction network has more impact than 3D-structure information.

#### 3.3.2 Vectors representing proteins in interaction network

We evaluate protein vectors that capture their relatedness in a protein-protein interaction network. To be consistent with DeepGO, we use the same protein vectors in their paper as input features for our GOAT. These vectors are created following the method in [1]. In brief, DeepGO uses vectors representing the protein names in a protein-protein interaction (PPI) network that has 8,478,935 proteins, and 11,586,695,610 edges total (derived from STRING database). Following [1], DeepGO uses DeepWalk [**?**] to generate sentences from the network, and apply Word2Vec [9] to these sentences to create the vector embedding for the protein names. Effectively, interacting proteins will have similar vectors.

Because Domain information is a key input of GOAT, we integrate PPI network data on top of our GOAT_MOTIF_. GOAT_MOTIF,PPI_ exceeds the other Transformer models by large margins (Table 1 row 8). On the entire label sets, in the presence of PPI network information GOAT and DeepGO perform similarly, despite the fact that GOAT can better extract information from the amino acid sequence (Table 1 row 4 and 8). Only for rare MF labels does GOAT exceed DeepGO at every R@k by noticeable margins (Table 2 row 4 and 8).

Intuitively, it is reasonable that PPI network dominates the information from amino acid sequences at classifying BP and CC labels, but not for MF labels. For example, two interacting proteins can have distinct 3D structures and sequences (and thus motifs); yet, they involve in the same biological process and sometimes found at the same cellular components. The same two interacting proteins however can have dissimilar molecular functions because they can induce very different chemical reactions.

We emphasize that vectors of PPI network in DeepGO does not need amino acid sequence to retrieve vectors representing the proteins. However, this method will not return vector representations for novel proteins not yet existed in the database. In practice, for proteins not yet well studied, we may need a different approach and other metadata for such proteins. We reserve this topic for future research work.

### 3.4 Evaluation on sparse GO labels

Many GO terms annotate only a few proteins because protein functions can be very unique. In practice, parametric predictors must handle sparse labels to predict terms that closely resemble the true protein functions. Yet, parametric models can fail when the train data has too many sparse labels. In such cases, GOAT and DeepGO accuracy will drop, because we need a label to have enough samples to reliably train the parameters in a neural network model. It is then important to evaluate GOAT and DeepGO against the nonparametric MetaGO_BLAST_ which does not have trainable parameters.

We evaluate GOAT and DeepGO on datasets that contain more rare labels. We reuse the same proteins in the original DeepGO data but include labels with at least 50, 10, and 10 occurrences in the BP, MF and CC train data, whereas the same criteria in the original DeepGO are 250, 50 and 50. Our larger datasets now have 2980 BP, 1697 MF and 989 CC labels, respectively (versus the original 932 BP, 589 MF, and 439 CC labels). For each added term, we include its ancestors as the gold-standard labels, so that most of the labels in the original data now have higher occurrence frequencies.

We train the models on the entire larger dataset. In this experiment, we also include a GOAT_PPI_ to evaluate the contribution of the PPI network data alone, and a GOAT_MOTIF,3D,PPI_ to evaluate the most complete model. In this experiment, we are less interested in the common labels, and evaluate R@k for the extra 2048 BP, 1108 MF and 550 CC labels which are sparse compared to the labels in the original DeepGO data; for example, 95% of the extra labels occur below 252, 34, and 68 times in the BP, MF and CC train data, respectively. Table 3 shows the R@k for these sparse labels.

**Table 3:** We increase the label size in the original DeepGO data from 932 BP, 589 MF, and 439 CC labels to 2980 BP, 1697 MF and 989 CC labels. Models are trained on the entire label sets, but R@k are evaluated only for the added 2048 BP, 1108 MF and 550 CC labels which are sparse. R@k are computed with respect to only proteins having these labels; there are 7850, 2671, and 1848 such proteins out of 9095, 6294, and 8886 samples in BP, MF and CC test data.

|  |  | BP |  |  | MF |  |  | CC |  |  |
| --- | --- | --- | --- | --- | --- | --- | --- | --- | --- | --- |
|  |  | R@40 | R@70 | R@100 | R@40 | R@70 | R@100 | R@10 | R@30 | R@50 |
| **BLAST Psi-BLAST** |  |  |  |  |  |  |  |  |  |  |
| 1 | Evalue10 | 30.30 | 34.84 | 37.88 | 38.67 | 41.09 | 43.42 | 20.80 | 29.15 | 33.13 |
| 2 | Evalue100 | 29.53 | 34.57 | 38.02 | 38.58 | 43.32 | 45.81 | 23.47 | 35.23 | 39.78 |
| **DeepGO** |  |  |  |  |  |  |  |  |  |  |
| 3 | BASE | 23.14 | 27.98 | 32.01 | 16.62 | 23.83 | 29.60 | 22.49 | 31.53 | 37.96 |
| 4 | +PPI | **48.06** | 55.86 | 61.02 | 41.61 | 48.61 | 53.39 | 47.89 | 60.77 | 66.48 |
| **GOAT** |  |  |  |  |  |  |  |  |  |  |
| 5 | BASE | 23.78 | 28.82 | 32.69 | 19.41 | 27.88 | 34.19 | 25.22 | 35.98 | 42.53 |
| 6 | +MOTIF+3D | 27.70 | 33.50 | 38.03 | 30.52 | 38.74 | 45.44 | 25.73 | 37.82 | 44.42 |
| 7 | +PPI | 43.51 | 52.52 | 58.54 | 41.77 | 52.46 | 58.51 | 41.98 | 60.02 | 68.70 |
| 8 | +MOTIF+PPI | 46.22 | 55.06 | 61.08 | 45.69 | 55.32 | 61.11 | **50.08** | **67.15** | 73.89 |
| 9 | +MOTIF+3D+PPI | 46.78 | **56.24** | **62.49** | **50.92** | **60.77** | **66.64** | 47.96 | 65.30 | **74.36** |

In Table 3, the base GOAT and DeepGO perform worst than MetaGO_BLAST_. We emphasize that in this case, GOAT is still has better performance than DeepGO for MF and CC data, but not for BP data (row 3 and 5). The PPI network data appears to be the most important factor; for example, GOAT_MOTIF,3D_ is about the same as MetaGO_BLAST_, whereas all the models with PPI network achieve higher R@k than MetaGO_BLAST_ (Table 3).

When predicting biological processes, in the presence of PPI network embedding, the other protein metadata and the types of neural network model for amino acid sequences are not as important (Table 3 row 4, and 7–9). This result is an empirical evidence supporting our earlier hypothesis that PPI network can dominate sequence information; that is, two interacting proteins should be involved in the same biological processes even when their sequences display dissimilar motifs, 3D structures, or any other types of hidden information to be extracted by neural network models.

For MF and CC labels, integrating just PPI network embedding into DeepGO is not enough to raise recall rates; in this case, our GOAT model, Domain information and SCOP data are important complementary factors to the PPI network embedding. Indeed, for MF labels, in the presence of PPI network embedding, GOAT also benefits when having Domain and 3D-structure information as extra features (Table 3 row 7 and 9). For CC labels, the same observation holds true, except that Domain information now has more impact than 3D-structure data (Table 3 row 8 and 9).

## 4 Discussion

In this paper, we introduce the novel **GO a**nnotation method with **T**ransformer (GOAT). We show that for predicting protein annotations, our Transformer architecture in GOAT is better than the convolutional neural network in DeepGO. We then provide GOAT three types of extra features: Domain information, 3D-structure and PPI network data. These features further increase the accuracy of GOAT, but PPI network information has the most impact.

Previous software MetaGO of Zhang <u>et al.</u> [18] has also combined sequence data, 3D-structure and PPI network information to annotate GO labels. We emphasize that in MetaGO, each type of metadata is used to build its own classifier, and then these independent classifiers are then combined to produce the final prediction for a GO label and an input sequence. For example, Zhang <u>et al.</u> [18] built their MetaGO_BLAST_ as an independent unit from the their two classifiers that uses PPI network and 3D-structure data. The reason for their strategy is that BLAST algorithm, which is similar to Smith-Waterman, does not need the interacting partners and 3D-structure of the input sequences [2]. Unlike BLAST-based methods, DeepGO and GOAT can jointly analyze the amino acid sequences, PPI network and 3D-structure information. In the future work, we wish to integrate components of MetaGO into GOAT, and vice versa.

We discuss a key property of our Transformer in the context of GO embeddings. This Transformer learns the co-occurrences among the labels; for example, the last layer in Transformer returns comparable vectors for a child and parent GO label (Fig. 1). GO embeddings produced by our Transformer are not equivalent to the embeddings produced by factorizing the co-occurrence matrix of GO labels, because GO embeddings from our Transformer are also affected by information from the amino acid sequence (Fig. 2). For our future work, we will integrate embedding learned from co-occurrence frequencies into our Transformer framework.

We outline a two key limitations of our adaptation of Transformer. First, in this paper, to make our software GOAT accessible to many users, we have reduced the standard number of parameters that Transformer often assume in other machine learning applications; for example, Rives <u>et al.</u> [12] trained a language model on protein sequences using a 36-layer Transformer. We expect that our GO annotation accuracy to increase if we train our model with more parameters and on more samples from the Uniprot database.

Second, we do not pre-train our Transformer. For example, before predicting GO labels, Transformer can be trained only on protein sequences with the following objective. We can remove amino acids from a sequence, and then use Transformer to retrieve these missing amino acids. Pre-training helps the parameters in Transformer to converge better for the latter tasks; however pre-training requires a lot of data, for example Rives <u>et al.</u> [12] pre-trained their model on 250 million sequences. For our future work, we will consider training a large-scale Transformer model to predict GO labels for protein sequences.

## 5 Appendix

**Figure 3:**
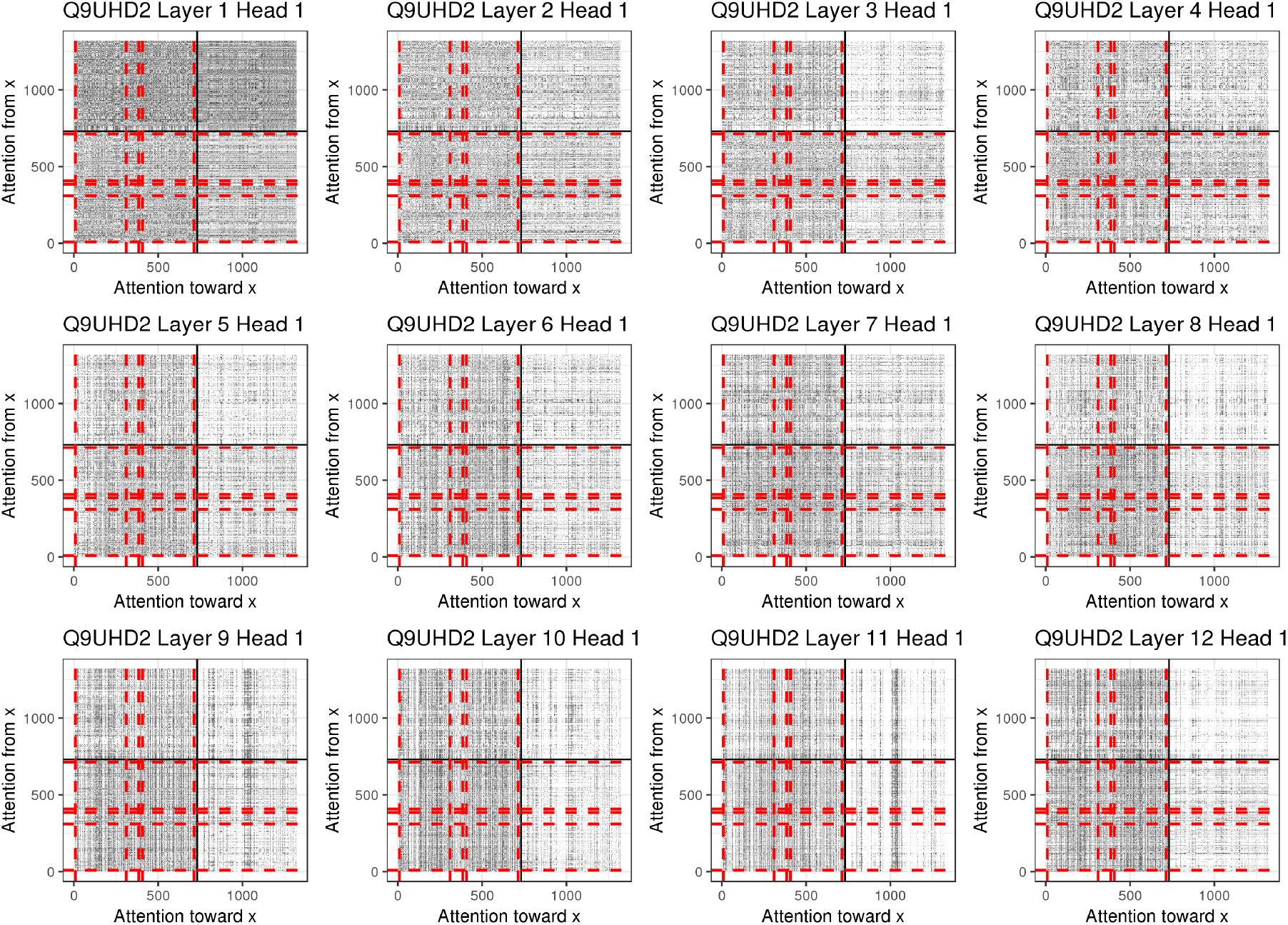
Heatmap of the attention values *α*_*jk*_ in each layer. Motifs of the sequences are not explicitly given as inputs to this Transformer model.

## Notes

https://github.com/datduong/GOAnnotationTransformer

